# Sweet science: Exploring the impact of fructose and glucose on brown adipocyte differentiation using optical diffraction tomography

**DOI:** 10.1101/2024.09.04.611269

**Authors:** Pooja Anantha, Xiangdong Wu, Salaheldeen Elsaid, Piyush Raj, Ishan Barman, Sui Seng Tee

## Abstract

The thermogenic capacity of brown adipose tissue (BAT) has garnered much attention for its potential to regulate systemic energy balance. BAT depot size and function need to be tightly to prevent loss of metabolic homeostasis due to energy dissipation via non-shivering thermogenesis. While adipocyte-intrinsic mechanisms controlling thermogenesis are critical, an increasing appreciation for the role of the BAT microenvironment is emerging. For example, changes in circulating hexoses due to dietary intake have shown to impact BAT function. Here, we show that murine BAT preadipocytes metabolism is impacted when fructose is used as the sole carbon source. Similarly differentiation medium containing only fructose yield mature adipocytes with fewer lipid droplets, with a concomitant decrease in adipogenic genes. These deficiencies are also observed in human BAT preadipocytes, where cutting-edge optical imaging modalities show a decrease in total cell mass and lipid mass in fructose-only medium. Taken together, the metabolic microenvironment significantly impacts BAT growth and function, with implications for the role of diets potentially mitigating the efficacy of BAT-targeted therapies.

## 1. INTRODUCTION

While white adipose tissue (WAT) is classically seen to store energy, BAT uniquely dissipates energy as heat. This is facilitated by uncoupling respiration from ATP synthesis, mediated through uncoupling protein (UCP)-1, located on the inner membrane of the mitochondria^1^. BAT is mainly found in the supraclavicular region, abundant in newborns as they lack the ability to shiver^2^. This thermogenic capacity has elicited much interest, with BAT described as a ‘metabolic sink’ that is associated with improved metabolic and cardiovascular outcomes^3^. Beyond its thermogenic capacity, BAT has also been described as an endocrine organ, secreting ‘batokines’ that promote improved metabolic health^4^.

Strategies to activate BAT thermogenesis have been intensely studied, ranging from cold acclimation to pharmacological activation, to more recent reports of BAT transplantation in preclinical models. A 10-day cold acclimation protocol is sufficient to increase BAT activity and non-shivering thermogenesis (NST) in healthy human volunteers, in a study reported more than a decade ago^5^. Nevertheless, intense cold exposure is impractical clinically, and increases the risk of adverse cardiovascular events^6^, leading to the development of pharmacological strategies to activate BAT thermogenesis. As both BAT thermogenic capacity (BAT differentiation, proliferation and UCP1 content) and activity (uncoupling activity and lipolysis)^7^ are controlled by central nervous system (CNS) via sympathetic output innervation, sympathomimetic drugs have been a central focus of research. These include mirabregon, an FDA-approved β3-receptor agonist, recently reported to be associated with a doubling of BAT mass, increased energy expenditure and improved insulin sensitivty^8^ in healthy women. Finally, an increasing body of literature in animal models^9^ have reported improvements in body weight/composition^10^, energy expenditure^11^, glucose/insulin sensitivity^12^ and circulating cytokines^13^ upon BAT transplantation. Collectively, these strategies rely on the ability of experimental interventions to promote and sustain BAT activation, potentially in challenging microenvironments, over sustained periods of time.

Thus, the metabolic microenvironment of BAT may have an outsized impact on the efficacy of BAT-targeted therapies. Nutritional deficiencies and/or overabundance can have consequences on BAT capacity and activity. BAT originate from Myf5^+^ stem cells of the skeletal muscle lineage. These cells are exquisitely sensitive to changes in nutrient signaling – as evidenced by reduced differentiation of preadipocytes to BAT upon knockout of insulin receptor (InsR)^14^. Strikingly, muscle development appeared normal in these mice, suggesting BAT-specific impacts of InsR deficiency. More recently, two weeks of high fructose feeding was shown to impair BAT glucose uptake in humans^15^, as quantified by radioactive fluorine-18 (^18^F) fluorodeoxyglucose (FDG) positron emission tomography (PET) scans. FDG-PET has traditionally been used as a surrogate for BAT activation^16^, and impaired radiotracer uptake appeared independent of insulin, and also unrelated to alterations of the microbiome. This study is consistent with preclinical studies, where a 12-week regime of high-fructose feeding in mice led to BAT ‘whitening’, that is a decrease of thermogenic capacity^17^. At the molecular level, the impact of fructose on BAT function remains ill-defined. Experiments involving the use of fructose in cell culture have been limited to murine embryo-derived, 3T3-based cell lines that can differentiate to adipogenic fate^18^. In these cells, supplementation of fructose to differentiation medium promotes adipogenesis in a concentration-dependent manner, that can be reversed by knockdown of the facilitative glucose transporter (GLUT)-5^19^. A subsequent report confirmed these observations in the same cell line, and also showed enhanced lipolysis, as quantified by an increase of hormone sensitive lipase (HSL) and adipocyte triglyceride lipase (ATGL)^20^. However, whether these phenotypes extend to thermogenic BAT remains unstudied. Given the exponential increase in fructose consumption, especially in the form of sugar-sweetened beverages^21^, there is an urgent need to define the potential impacts of a microenvironment high in fructose on BAT capacity and activity.

Imaging platforms that can characterize BAT physiology with minimal physiological perturbation will be invaluable to understand the role of the metabolic microenvironment. Metabolic flux is rapid and dynamic; the ideal tool to exemplify BAT biology would be similarly fast, without the need to introduce external contrast agents that may impact metabolism. We have previously shown the potential for label-free approaches to reveal the dynamic changes in cellular phenotypes when adipocytes undergo directed differentiation. Optical diffraction tomography (ODT) exploits the differences in the phase shift of light waves passing through objects, to generate a 3D refractive index (RI) map of the cell^22-24^. ODT is a label-free technology that enables the detailed visualization of cellular structures without introducing external contrast agents, making it ideal for studying the dynamic changes in cell morphology and composition during differentiation.

In our study, we employed ODT to image human brown preadipocytes (HBP) at multiple time points (0, 7, 14, 21 days) during adipogenic differentiation. We exposed HBPs to varying conditions of fructose, glucose, and a combination of both sugars to observe their effects on cell morphology and lipid formation throughout the differentiation process. To the best of our knowledge, this is the first study to utilize ODT in such a manner, providing novel insights into how specific sugars influence BAT cell development. Our previous study^25^ marked the first use of ODT to study human brown adipocytes ever, and here we are expanding our knowledge by studying the effect of sugars on adipogenic differentiation. Taken together, the insight generated from these studies provide a platform to understand the consequence of sugars on BAT function, as well as providing tools to dynamically characterize BAT physiology, that can be translated in the future to assess strategies that promote BAT activity to combat metabolic disease. Moreover, this work is pioneering in quantifying key cellular metrics, such as HBP cell mass, cell volume, lipid volume, and lipid diameter, under these conditions—calculations made possible through the precise imaging capabilities of ODT offering picogram precision. These measurements are critical for understanding how different sugar environments impact the physical properties of differentiating brown adipocytes, offering new perspectives on the role of the metabolic microenvironment in BAT function.

## 2. METHODS

### Isolation and immortalization of murine stromal vascular cells

Brown fat tissue was removed from interscapular region, minced, digested (1 mg/ml collagenase II at 37C for 30-40 min) and filtered through a 100μm cell strainer. Cells were then plated into a 6-well culture plate with DMEM/F12, 10% FBS, and 1% penicillin-streptomycin in 37C and 5% CO2 and immortalized with retrovirus expressing SV40 Large T antigen. Cells were cultured with neomycin selection for at least 7 days.

### Differentiation of mouse primary brown adipocyte

Immortalized preadipocytes were grown in the DMEM supplemented with 1g/L glucose, 10% FBS and 1% penicillin-streptomycin. When cells reached to 100% confluence, differentiation (day 0) was initiated with the induction medium with 1g/L glucose or 1g/L fructose supplemented with 0.5mM IBMX, 20 nM insulin, 125nM indomethacin, 1nM T3, 1μM dexamethasone, 1 μM rosiglitazone for 48 hours. Cells were then incubated in maintenance medium with 1g/L glucose or 1g/L fructose supplemented with 20nM insulin and 1nM T3.

### Lipid droplet staining

Live cell lipid droplet staining was achieved by HCS LipidTOX deep red neutral lipid stain and imaged with fluorescence microscopy (Leica DMi8 system).

### Culture and differentiation of human brown preadipocytes

Immortalized human brown preadipocytes (HBPs) were originally isolated by the Tseng Lab from the stromal vascular fraction of human adipose tissues, and then immortalized with hTERT^26-28^. We followed the published protocol to maintain and differentiate the HBPs^27,28^. HBPs were cultured and maintained in high-glucose Dulbecco’s modified Eagle’s medium (DMEM/H, Sigma-Aldrich) supplemented with 10% fetal bovine serum (Corning Cellgro), 100U/ml penicillin, and 100U/ml streptomycin (Thermofisher), in a humidified incubator in 5% CO_2_ at 37 °C. Cells were subcultured when they reached 90% confluence. To initiate adipogenic differentiation, cells were first grown in culture medium until confluent, and then switched to induction medium (IM), which is culture medium supplemented with 33 μM biotin, 0.5 μM human insulin, 17 μM pantothenate, 0.1 μM dexamethasone, 2 nM T3, 500 μM IBMX, and 30 μM indomethacin. All the ingredients for induction medium were purchased from Sigma Aldrich. Culture medium was changed once every 72 hrs.

To study the effects of fructose and glucose on adipogenic differentiation, the following conditions were studied:

Condition 1 (C1): IM with high glucose (25mM), serving as control.

Condition 2 (C2): IM with no sugar, serving as a second control.

Condition 3 (C3): IM with fructose (5mM).

Condition 4 (C4): IM with fructose (5mM) and glucose (5mM).

Condition 5 (C5): IM with fructose (5mM) and glucose (15mM).

HBPs were exposed to above conditions on “day 0”. “Day 1” marks the first day that the HBPs are exposed to the above conditions for a full 24 hours. Media was changed once every 72 hrs.

### Optical Diffraction Tomography and Analysis

HBPs were seeded on glass-bottom dishes (TomoDish) and fixed with 4% paraformaldehyde (PFA) 0, 7, 14, or 21 days after the initiation of differentiation (referred to as “day 0”, “day 7”, “day 14”, and “day 21” cells). Cells from the same batch were used to prepare all the samples to minimize variations between batches. After fixation, images of cells were captured using a holotomography system (HT-2H, TomoCube, Republic of Korea) comprised of a motorized stage, a 60x, 1.2 NA water-immersion objective, a 532 nm laser, and a dynamic micromirror device, that measure the 3D refractive indexes. TomoStudio (TomoCube, Republic of Korea) was used to visualize and obtain 3D RI tomogram images and 2D maximum intensity projection (MIP) images. Image processing was done using ImageJ. Cell segmentation was performed manually on 2D MIP images using ImageJ. The 2D segmented cells were used as masks and run through the stacks of 3D RI tomogram images to obtain 3D RI tomogram images with only one cell per field of view, as multiple cells are usually present in the original 3D RI tomogram images. Quantitative analyses were performed on 3D RI tomogram images with one cell per field of view using MATLAB. Cell dry mass and lipid mass values were obtained by initially setting refractive index thresholds and then calculating the surface integral of the optical path difference over the specific RI increment^23^. Cell volume and lipid volume were calculated by calculating the total number of bright pixels based on RI. Lipid diameter was calculated by assuming the LD was a sphere and obtaining the diameter accordingly.

## 3. RESULTS and DISCUSSION

### 3.1. Hexose availability impacts murine brown preadipocyte growth and metabolism

Optimal BAT function requires both preadipocyte hyperplasia (cell number increase) and hypertrophy (cell size increase)^29^. Thus, we first evaluated the ability of murine brown preadipocyte (MBP) proliferation, in an undifferentiated state, in the presence of different hexose microenvironments.

Conventional brown adipose growth medium utilized glucose at high concentrations (25 mM). Here, we elected to use glucose at 5 mM, to mirror physiological glucose concentrations, as the baseline condition to which other hexose conditions can be compared. To our surprise, growth medium with no hexose supported preadipocyte growth, albeit at a slower rate than glucose as the sole carbon source. In contrast, replacing glucose with equimolar fructose did not impact MBP proliferation. Similarly, a combination of 5 mM each of glucose and fructose or 10 mM glucose yielded similar growth rates as glucose alone (Fig. 1A and B).

**Figure 1:**
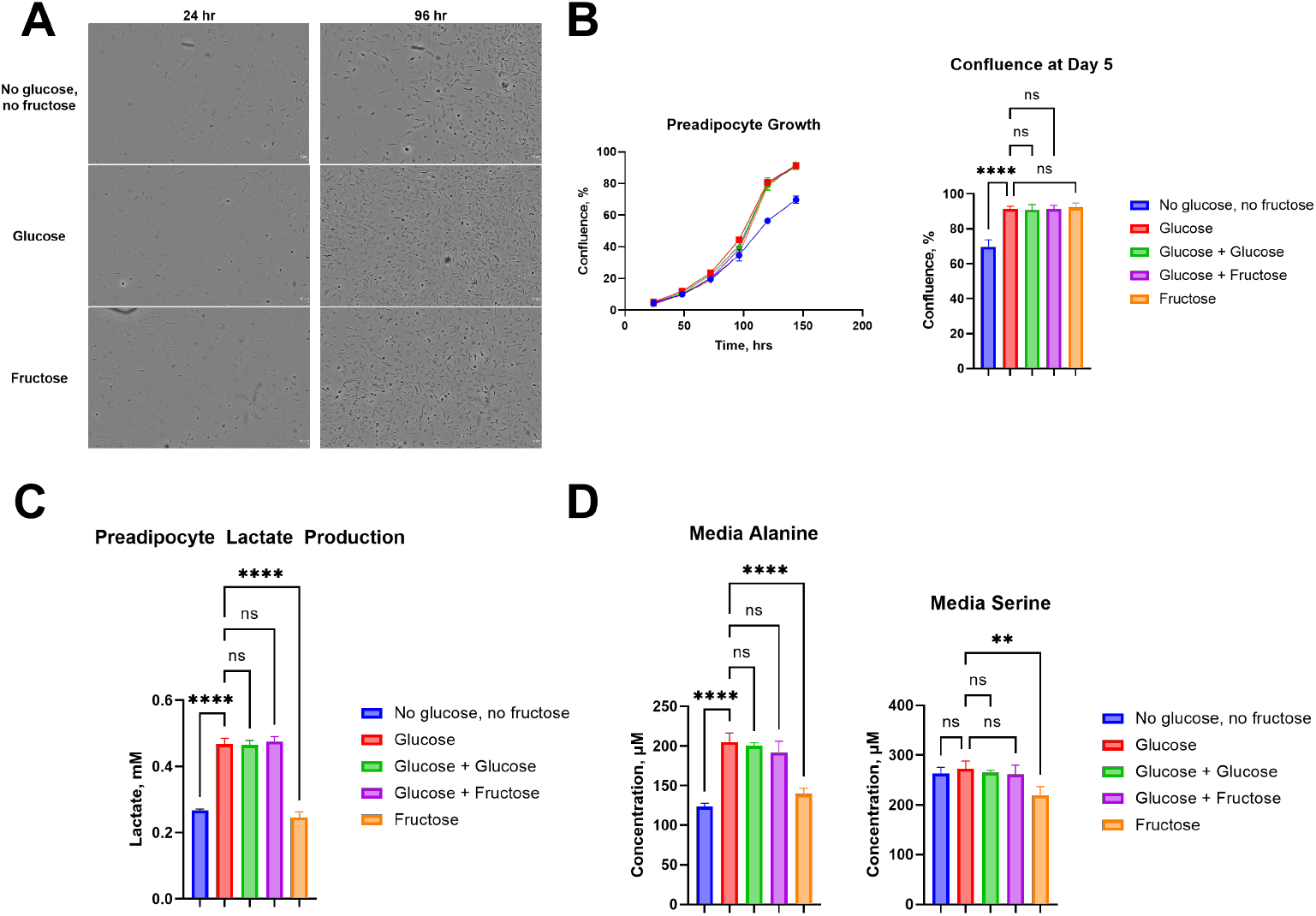
A) Representative images of MBP grown in either glucose or fructose as the sole carbon source, or in the absence of hexose. B) Growth curve of MBP quantified as cellular confluence, where MBP grown in the absence of hexose showing significantly reduced growth at day 5, compared to culture medium containing either individual or a mixture of hexoses (5mM of each hexose). C) Quantification of lactate and D) amino acid in spent growth medium of MBP cultured in different hexoses.

While fructose and glucose both supported MBP proliferation to similar degrees, we proceeded to ask if these hexoses differentially impacted cellular metabolism. Lactate is a common metabolic fate of both glucose and fructose oxidation, and we quantified lactate concentrations in MBP growth medium after 6 days of growth. Predictably, the absence of hexose resulted in the lowest lactate concentrations in these cells, possibly as a consequence of reduced proliferation. However, MBP grown in fructose alone also showed a significant decrease in lactate production, with ~50% decrease compared to cells grown in equimolar glucose.

Beyond hexoses, amino acids are also critical to maintain metabolic homeostasis during cell growth. Here, we observed reduced alanine in the spent medium of cells grown without hexoses, as well as in fructose alone, that is significantly different compared to cells grown in 5 mM glucose. Serine concentrations in spent medium of fructose-grown cells were also significantly reduced, but this effect is not seen in cells grown without hexoses.

Thus, while MBP demonstrated remarkable flexibility in hexose utilization to maintain proliferation, fructose reprograms both lactate and amino acid metabolism, relative to glucose.

### 3.2. Hexose availability impacts brown preadipocyte differentiation

We proceeded to evaluate the impact of hexoses on preadipocyte hypertrophy, by quantifying the ability of BAT preadipocyte to differentiate into mature adipocytes. This was enabled by the combined presence of insulin, dexamethasone and tri-iodothyronine (Fig. 2A).

**Figure 2:**
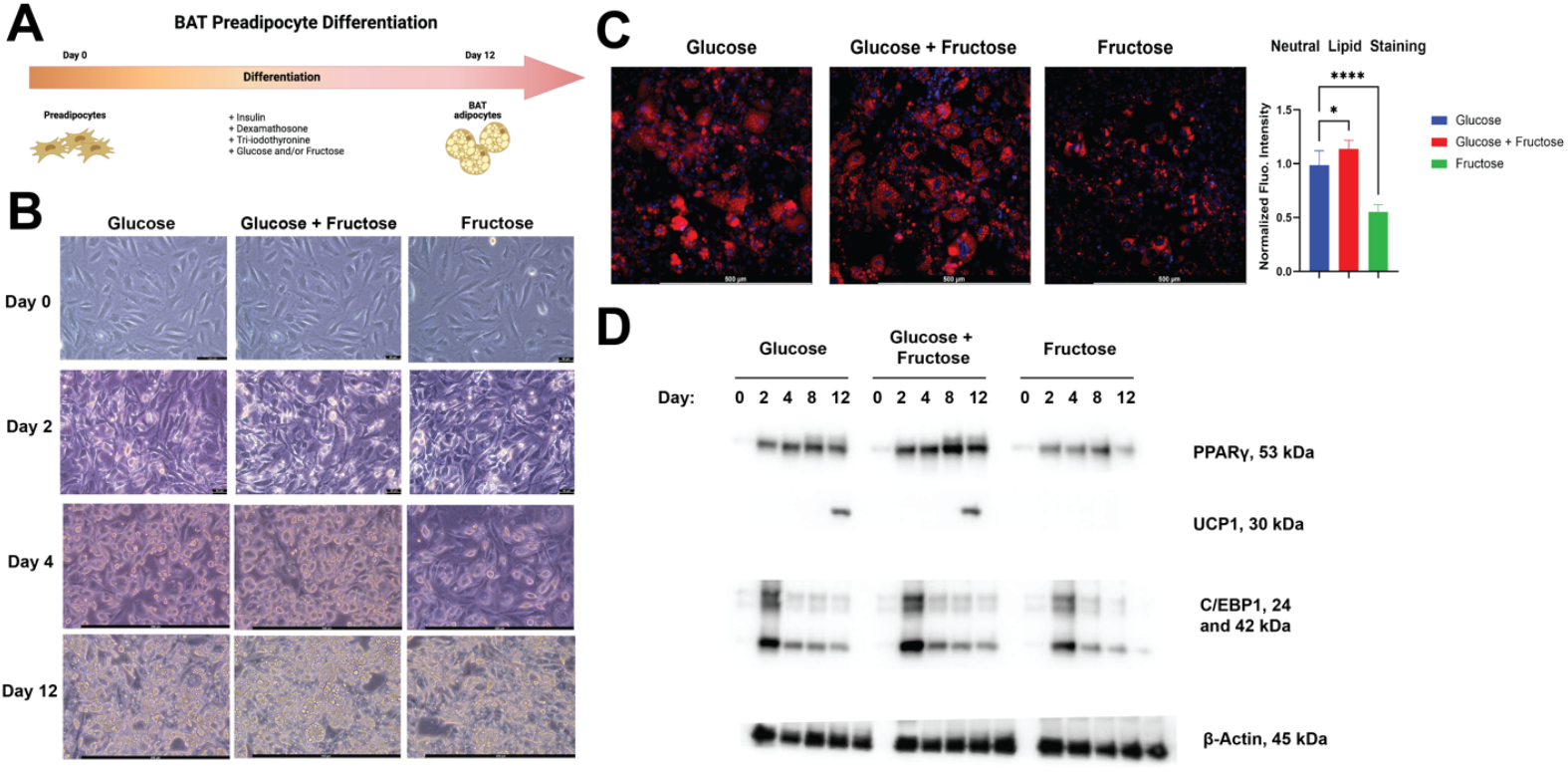
A) Schematic of MBP differentiation over a 12-day period. (B) Representative images of MBP over the course of 12 days, where glucose, fructose or a mixture resulted in striking morphological differences. (C) Fluorescence staining of intracellular neutral lipids, where MBP grown in fructose as the sole carbon source showed a significant reduction in lipid content. (D) Western blots of total cell lysate, showing reduced expression of critical regulators of BAT adipogenesis (PPARG and CEBP1) and thermogenesis (UCP1) in MBP grown in fructose as the sole carbon source.

Here, we compared the addition of fructose (5 mM), to standard differentiation medium containing basal levels of 5 mM glucose. In addition, we also evaluated the ability of 5 mM fructose as the sole carbon source to support adipogenic differentiation. Addition of these growth factors resulted in the formation of distinct lipid droplets in cells, that increased in a time-dependent fashion over a period of 12 days in both glucose only, and glucose + fructose culture medium. In contrast, fructose as the sole hexose resulted in an observable decrease of lipid droplets, evident as early as 4 days post-differentiation (Fig. 2B).

Confirmation that these organelles contained lipids were performed by using the hydrophobic BODIPY dye, that preferentially binds neutral lipids. Quantification of BODIPY fluorescence intensity, normalized by the nuclear stain DAPI, showed ~50% decrease in lipid content in cells cultured in fructose alone, compared to glucose alone. In contrast, the addition of fructose to glucose resulted in a significant increase in lipid content. Together, these results suggest that compared to glucose, fructose as the sole carbon source dampens adipogenesis. However, this defect can be reversed in the presence of glucose, and when both hexoses are present, adipogenesis is elevated compared to glucose as the sole substrate.

Beyond lipid droplet formation, differentiation of BAT preadipocyte to mature adipocyte is followed by concomitant upregulation of UCP1, that is responsible for thermogenesis. Immunoblotting for UCP1 levels in total cell lysates demonstrated robust expression of this protein 12 days after induction of differentiation in glucose-only medium. In contrast, fructose as the sole hexose resulted in the lowest levels of UCP1, while the presence of both hexoses coincided with the highest levels of this protein. UCP1 expression is regulated by the binding of transcription factors to the promoter regions, including binding sites for PPAR and C/EBP1. Accordingly, the levels of these transcriptional regulators were dampened in fructose as the sole hexose (Fig. 2D).

Taken together, preadipocyte hypertrophy via differentiation into mature BAT adipocyte is dampened when fructose is supplied as the sole hexose. This block in differentiation manifests phenotypically as a reduction in intracellular lipid droplets and UCP1 levels, both hallmarks of mature BAT.

### 3.3. Hexose availability impacts human brown preadipocyte (HBP) cell mass and volume

Thus far, we have shown that hexose availability significantly impacts MBP hyperplasia and hypertrophy, where media containing only fructose as a carbon source dampening adipogenesis. To evaluate the translational relevance of these observations, we extended our investigation to human brown preadipocytes (HBP), hypothesizing that the presence of different hexoses will also influence the growth and metabolism in human cells.

In this study, we utilized ODT to capture images of HBPs at days 0, 7, 14, and 21 of adipogenic differentiation under varying levels of hexose conditions (Figure 3A). Following imaging, we quantified several key parameters, including cell dry mass (Figures 3B-3I), cell volume (Figures 3J-3Q), lipid dry mass (Figures 4A-4H), and lipid volume (Figures 4I-4P). Cell dry mass and volume provide insights into lipid accumulation, a crucial aspect of adipocyte differentiation. Additionally, we calculated the contribution of lipid dry mass to total cell dry mass (Figures 4Q-4X) to further understand the impact of different metabolic environments on HBP differentiation.

**Figure 3:**
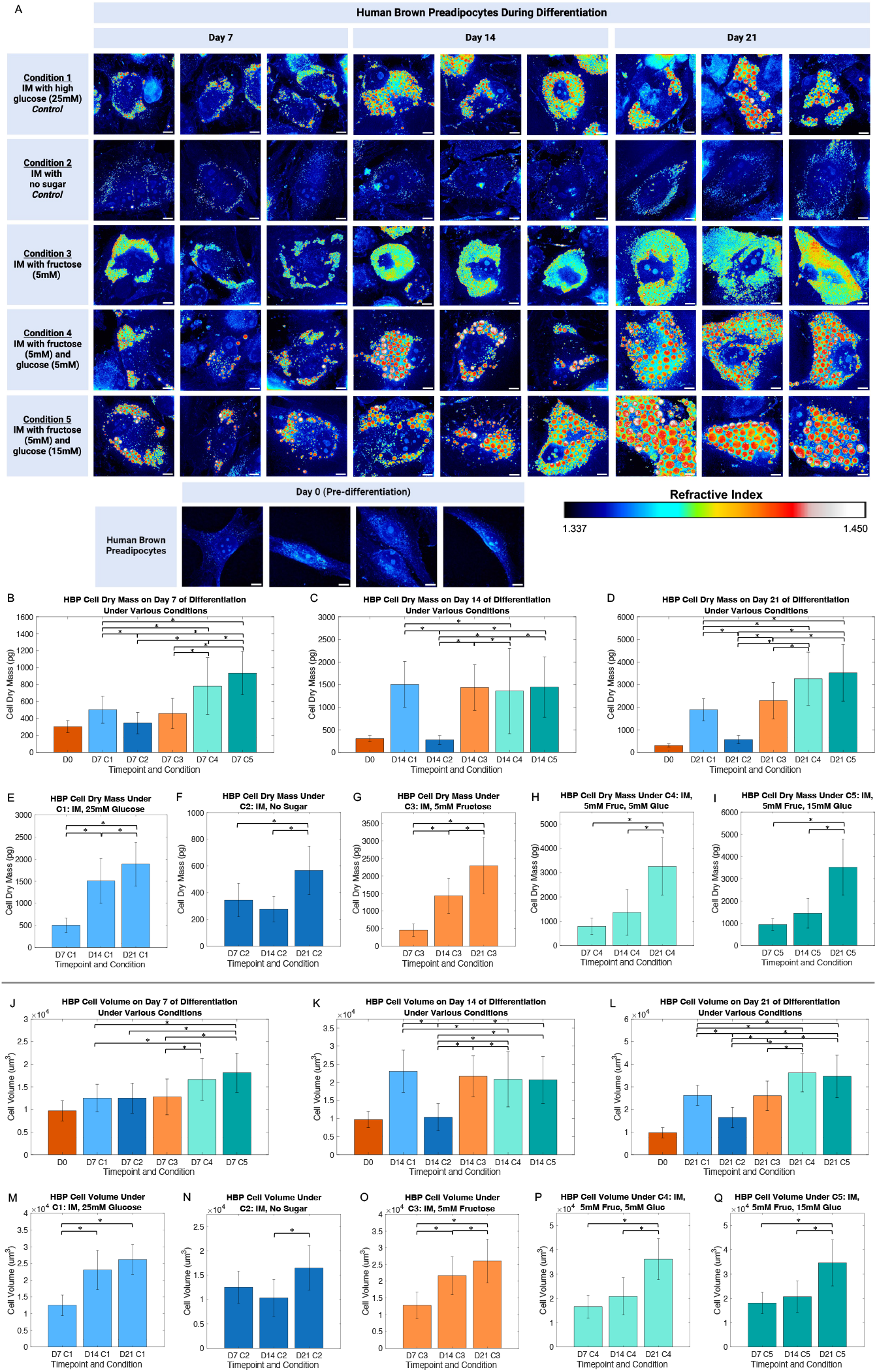
A) Optical diffraction tomography (ODT) images of human brown preadipocytes (HBPs) at day 0, 7, 14, and 21 of adipogenic differentiation under various conditions (scale bar: 10μm). Color bar indicates refractive index range. HBP cell dry mass on various days of differentiation under various conditions (B-I), HBP cell volume on various days of differentiation under various conditions (J-Q) (*p ≤ 0.05).

**Figure 4:**
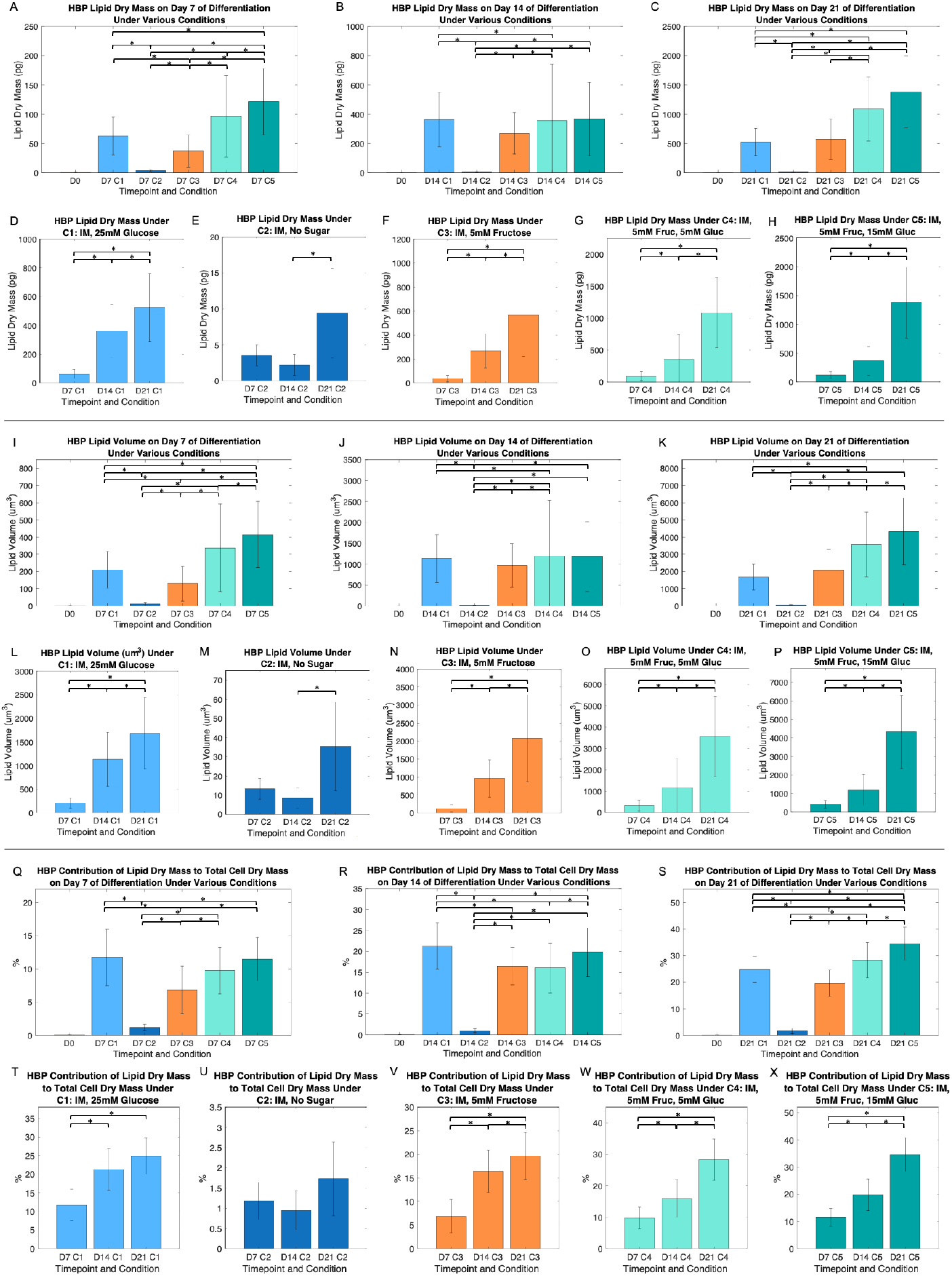
A) HBP lipid dry mass on various days of differentiation under various conditions (A-H), HBP lipid volume on various days of differentiation under various conditions (I-P), HBP contribution of lipid dry mass to cell dry mass on various days of differentiation under various conditions (Q-X) (*p ≤ 0.05).

The ODT images reveal that at day 0, preadipocytes exhibit minimal to no lipid accumulation, which is expected for undifferentiated cells. As differentiation progresses through days 7, 14, and 21, there is a noticeable increase in lipid accumulation across most conditions, demonstrating the effectiveness of the induction media in promoting adipogenesis. However, under condition 2 (C2), where no sugars are present in the induction medium, HBPs show little to no lipid formation at any time point. This indicates the essential role of sugars in enabling lipid droplet (LD) formation during adipogenesis. The absence of sugars in C2 leads to a failure in lipid accumulation, highlighting the necessity of hexoses for effective differentiation.

In contrast, condition 3 (C3), where fructose is only present, this results in the formation of smaller LDs compared to other conditions. This trend persists across all observed time points, suggesting that fructose alone may limit the expansion of lipid droplets during differentiation. On the other hand, the presence of glucose in condition 4 (C4) and condition 5 (C5) promotes the formation of larger LDs, indicating that glucose plays a significant role in lipid droplet size.

Figure 3B shows that on Day 7, there are significant differences between C1 and C2, with C5 exhibiting the highest mean cell dry mass of around 1000 pg compared to the other conditions for day 7. C2 is not significantly different from C3 but does differ from C4 and C5. On Day 14 (Figure 3C), cell dry mass increases overall. DC2 shows significant differences with all other conditions. The mean cell dry mass for C1, C3, C4, and C5 is around 1500 pg, with significant differences maintained among these groups. By day 21 (Figure 3D), the mean cell dry mass continues to rise, with C2 showing significant differences from all other conditions. C4 and C5 have similar mean cell dry masses of approximately 3500 pg and are significantly different from C3 and others, though not from each other.

Figure 3E highlights significant differences in cell dry mass for C1 across all three days. For cell dry mass due to C2 (Figure 3F), significant differences are found only between day 14 and day 21 and between day 7 and day 21, with no significant difference between day 7 and day 14. For cell dry mass due to C3 (Figure 3G) shows significant differences across all days. For cell dry mass due to C4 (Figure 3H) exhibits significant differences only between Day 14 and Day 21 and between day 7 and day 21, with no significant differences between day 7 and day 14. For cell dry mass due to C5 (Figure 3I), significant differences are observed between Day 14 and Day 21 and between day 7 and day 21, but not between day 7 and day 14. Figures 3H and 3I show that D7C4 and D14C4 are similar to D7C5 and D14C5, with mean cell dry masses of about 1000 and 1500 pg, respectively. D21C5 has the highest mean cell dry mass, slightly over 3500 pg. Overall, cell dry mass increases over time, with notable differences across conditions and days. C2 and C3 show consistent increases, while C4 and C5 exhibit significant growth by day 21.

Figure 3J shows cell volume on day 7 across different conditions. The mean cell volume for C1, C2, and C3 is similar, averaging around 12,500 μm^3^, and these values are not significantly different from each other. In contrast, C4 and C5 exhibit a noticeable increase in cell volume. On day 14 (Figure 3K), the mean cell volume for C1, C3, C4, and C5 ranges between 20,000 and 25,000 μm^3^. There are some significant differences among these conditions, but overall, the cell volumes are similar across C1, C3, and C5. This suggests that by day 14, conditions that initially led to increased cell volume (like C4 and C5) continue to support substantial cell expansion. By day 21 (Figure 3L), the mean cell volume reaches its peak, with C4 and C5 showing the highest volumes of approximately 35,000 μm^3^. These volumes are not significantly different from each other.

Taken together, hexose availability is necessary for HBP hypertrophy in the presence of adipogenic differentiation cues. However, the extent of adipocyte hypertrophy is highly heterogenous, where an increase of cellular volume is not linearly correlated to an increase in extracellular hexose concentrations. In contrast, the type of hexose significantly alters cell mass and volume, where the simultaneous presence of glucose and fructose results in cells with the greatest mass and volume, compared to those grown in glucose alone.

### 3.4. Hexose availability impacts human brown preadipocyte (HBP) lipid mass and volume

Expanding on our observation of altered cellular mass and volume, we proceeded to evaluate the contribution of lipids to changes in the biophysical properties of HBP. This is motivated by the heterogeneous nature of hexose metabolism, where the metabolic fate of sugars can be diverse, contributing to the biosynthesis of various macromolecules such as nucleic acid, proteins and lipids. Here, we utilize ODT as a minimally invasive tool, to provide critical insights into the effects of different hexose environments on HBP lipid content.

The negligible lipid presence at day 0 across all conditions aligns with the expected absence of adipogenic differentiation at this initial stage. This serves as a baseline, reinforcing that any subsequent lipid accumulation is a direct result of the differentiation process influenced by the specific conditions applied. For condition C2, the persistently low lipid dry mass across all timepoints is particularly noteworthy. This suggests that the specific metabolic environment in C2 is not conducive to significant lipid accumulation, indicating a potential inhibitory effect on adipogenesis. This could have implications for understanding how certain conditions or nutrients might suppress lipid formation in brown preadipocytes, which could be relevant in therapeutic strategies aimed at regulating fat accumulation.

The observed doubling of mean lipid mass in conditions C1 and C5 at all timepoints highlights the pronounced effect these environments have on promoting lipid accumulation. The highest mean lipid dry mass at day 21 under condition C5, reaching approximately 1500 pg, underscores the significant lipid synthesis and storage capacity of HBPs in this condition. This finding is critical as it suggests that certain metabolic environments, particularly those similar to C5, could enhance lipid accumulation, potentially influencing the energy storage and thermogenic capabilities of brown adipocytes. Similarly, the doubling trend observed in lipid volume from day 7 to day 21 in conditions C3-C5 further emphasizes the impact of these conditions on HBP development. The highest mean lipid volume recorded at day 21 under condition C5, around 4500 μm^3^, indicates that not only is the lipid mass increasing, but the physical space occupied by lipids within the cells is also expanding significantly. This could have implications for the overall morphology and functionality of brown adipocytes, as larger lipid volumes might correlate with enhanced thermogenic potential.

We also examined how lipid dry mass contributed to the total cell dry mass across various timepoints and conditions. This analysis provided further insights into the extent of lipid accumulation relative to the overall cell composition during differentiation. For all timepoints, condition C2 exhibited a consistently low contribution of lipid mass to total cell dry mass, accounting for less than 5%. This finding aligns with our earlier observations from the ODT images, where C2 showed minimal lipid accumulation, reinforcing the notion that this specific condition is not favorable for adipogenic differentiation in HBPs. In contrast, condition C3 demonstrated a more substantial contribution, with about 20% of the total cell dry mass attributed to lipid mass. This indicates that C3 supports a moderate level of lipid accumulation during differentiation. More pronounced effects were observed in conditions C4 and C5, where lipid mass contributed to 30% and 40% of the total cell dry mass, respectively. These findings suggest that the metabolic environments in C4 and C5 are particularly conducive to lipid synthesis and storage within HBPs, leading to a significant proportion of the cell’s dry mass being composed of lipids. In addition, our current study results for HBP cell dry mass and lipid dry mass are consistent with the results of the study reported previously^25^.

### 3.5. The type of hexose affects HBP lipid size and quantity during adipogenic differentiation

Hexose availability has striking effects on cell size and volume as HBP differentiate, with intracellular lipid content contributing significantly to this change. We wanted to understand the heterogeneity of these lipids, as a function of the type of hexose available to cells.

Figure 5 shows the lipid area contribution (%) versus lipid diameter distribution —categorized by timepoint and condition. This analysis provides a more detailed understanding of how different sugars influence the quantity and size distribution of lipid droplets during differentiation. Under C3 across all timepoints, the highest percentage of lipid area contribution consistently came from lipid droplets (LDs) smaller than 1 μm, indicating that HBP cells exposed to fructose alone tend to form the smallest LDs compared to any other condition at any timepoint. As the conditions progressed from C3 to C4 to C5 across all timepoints, there was a notable increase in the percentage of lipid area contribution from LDs with progressively larger diameters. This trend suggests that the combination of fructose with increasing concentrations of glucose results in the formation of more and larger LDs within HBP cells.

**Figure 5:**
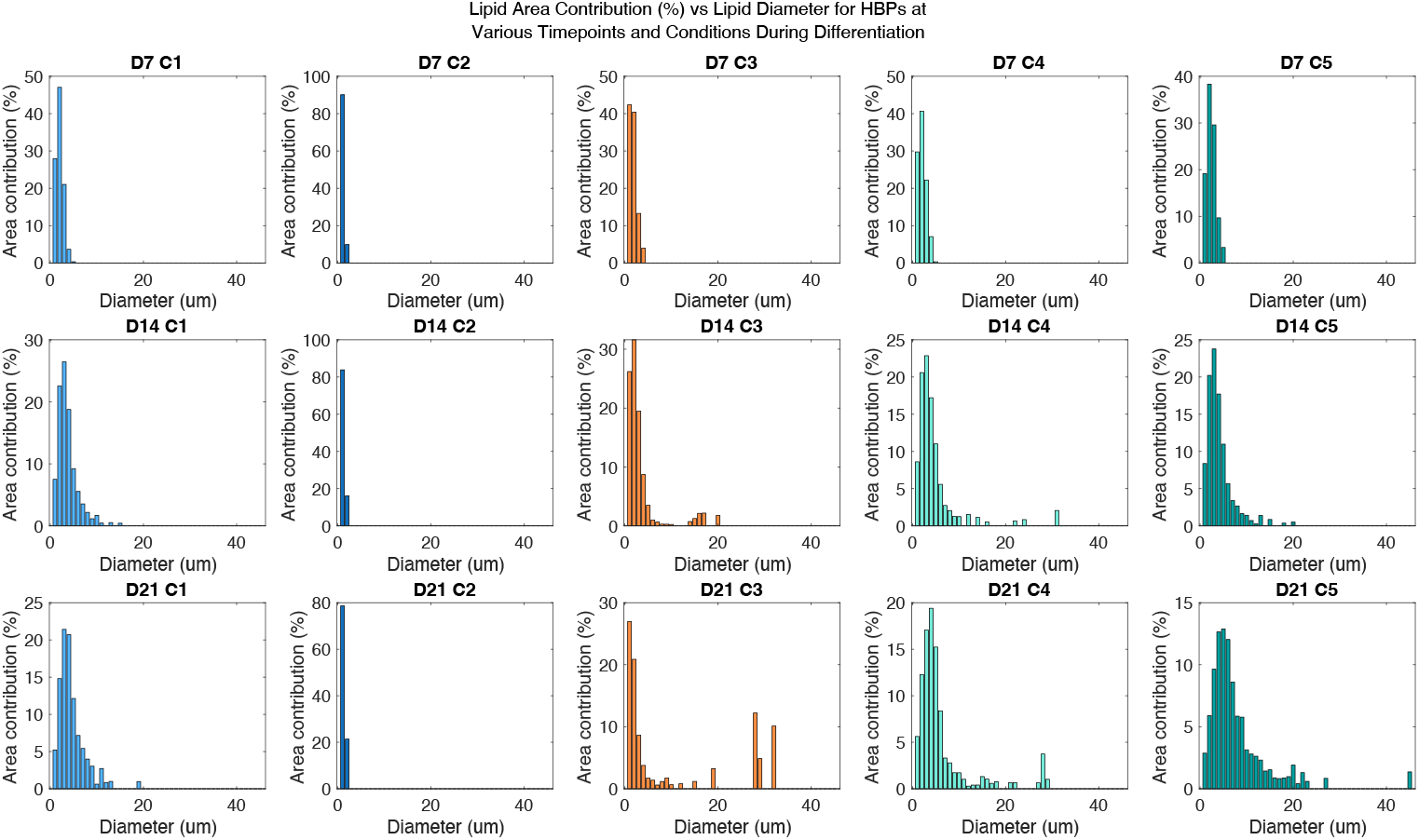
Lipid area contribution (%) vs lipid diameter distribution for HBPs at various timepoints and conditions during adipogenic differentiation.

Furthermore, when comparing day 21 to day 7 for conditions C1, C3, C4, and C5, we observed a marked increase in the percentage of lipid area contribution from LDs on Day 21, reflecting the presence of more and larger LDs as HBP differentiation progresses over time. In contrast, under C2 across all timepoints, nearly all lipid area contribution came from LDs smaller than 1 μm, with the total lipid mass comprising less than 2% of the cell dry mass. This outcome indicates that in the absence of sugar, there is minimal to no formation of LDs, despite the presence of adipogenic differentiation medium. The presence of large LDs, particularly the 30 μm LDs observed in day 21 under C3, represents a notable discrepancy. These unusually large LDs are not simply individual droplets but are the result of smaller LDs fusing together, which is supported by their irregular shapes rather than the typically spherical morphology of single LDs. This phenomenon of fused LDs accounts for less than 2% of all LDs observed in our cells. Additionally, we identified a few other large LDs in day 21 under C4 and C5, including one nearly 45 μm in diameter. The diameters of fused LDs were calculated using a best-fit ellipse and taking an average of the major and minor axis. An example of these irregularly shaped, fused LDs is shown in Figure S1. These findings provide critical insights into how specific sugar environments influence the lipid characteristics of HBP cells during adipogenic differentiation. Thus, directed differentiation of HBP in the presence of different types of hexose results in heterogeneous lipid phenotypes, with striking differences in total lipid content and lipid droplet diameters.

## DISCUSSION

BAT is best known for thermogenesis by increasing whole-body energy expenditure. However, the nature of the relationship between ingested nutrients and BAT function remains incompletely defined. While fructose has been associated with hyperlipidemia^30^, hyperinsulinemia^31^, and obesity^32^, a causal role of fructose in promoting BAT dysfunction has not been established. However, two recent studies have pointed to fructose negatively impacting BAT function. The first study in rodents show that fructose increases intestinal acetate production, mediated by the microbiome^33^. As direct injection of acetate into BAT inhibits thermogenesis^34^, it is possible that fructose-induced, microbiome-derived acetate may impair BAT physiology. A more direct role for fructose has also been implied, where 10 healthy, young (20 – 40 years), non-obese men were exposed to 2 weeks of high-glucose or high-fructose feeding. Strikingly, only fructose-feeding resulted in dampened glucose uptake by BAT, with no changes in blood biochemistry or microbial content. Thus, these lines of evidence support a potentially complex role of extracellular fructose on BAT physiology and function.

To clarify the role of fructose and related hexoses, this study utilized murine and human brown preadipocytes (MBP and HBP) as models for BAT growth and function. These preadipocytes are able to respond to differentiation cues, forming mature BAT with characteristic lipid droplets and expression of BAT-associated UCP-1^28^. Biochemical and biophysical characterization of MBP and HBP revealed significant divergence in both preadipocyte growth, adipogenesis and expression of thermogenesis related UCP-1 when these cells are exposed to fructose and/or glucose. The hyperplastic ability of MBP was retained, regardless of the type of hexose present – these cells proliferated at the same rate in either fructose or glucose, or both. Reduced MBP proliferation was only seen when cells were deprived of hexose. As preadipocyte hyperplasia is a contributor to cold adaptation mediated by BAT^35,36^, our experiments suggest that hexoses play a minor role in influencing BAT function, downstream of preadipocyte proliferation. While proliferation rates are unchanged, amino acid metabolism was altered in MBP. Alanine and serine levels in spent growth medium were lowered in fructose grown cells, compared to equimolar glucose. This could indicate either reduced production or increased consumption of these amino acids, that remain an interesting avenue for future studies. Of note, recent reports of BAT being a ‘metabolic sink’ for branched chain amino acids^37-39^ add significance to our observations, where hexose availability may act as a regulator of amino acid catabolism in thermogenic tissues.

In contrast to proliferation, hexose availability resulted in significant differences in preadipocyte differentiation. MBPs showed ~50% less neutral lipid accumulation in fructose compared to cells grown in equivalent amounts of glucose. Reduced adipogenesis is attributed to lower levels of two transcription factors, PPARG and CEBPa. Supporting this observation is the reduced levels of UCP-1, as the proximal region of the UCP-1 gene contains binding sites for CEBPa, that is augmented by binding of PPARG at a distal enhancer region^40^. Together, these observations suggest a mechanism where fructose imposes a block on adipogenic and thermogenic gene transcription via reduced expression of PPARG and CEBPa.

As different hexoses affected proliferation and differentiation in murine cells, we asked if similar alterations were observed in human brown preadipocytes (HBP). Here, we utilized biophysical methods to complement biochemical assessment of preadipocyte differentiation. Conventional HBP differentiation protocols utilizing 25mM glucose resulted in > 10X increase in cell dry mass and ~3X increase in total cell volume, along with the formation of numerous lipid droplets after 21 days. These dramatic phenotypes were lost when hexose was removed, suggesting that anabolic conversion of glucose-derived carbons into lipids account for most of the increases seen in mass and volume. Combinations of different hexoses also produced remarkably distinct changes in HBP biophysical properties. While circulating fructose levels are unlikely to exceed 5mM, serum glucose levels may vary, providing the rationale for conditions C4 and C5 used in HBP experiments, mirroring normo- and hyperglycemia. Relative to 25mM glucose, combinations of 5mM fructose with either 5- or 15mM glucose (C4 and C5, respectively) both resulted in cells with significantly larger mass and volume at 21 days. This was unexpected, as C4 and C5 contained fewer total carbons from hexose (10- and 20mM, respectively), relative to C1 (25mM glucose). Thus, our experiments suggest that increases in cell mass and volume during adipogenic differentiation is not a linear function of hexose carbon availability, highlighting the complex relationship between metabolism and cell size.

Replacing 25mM glucose with 5mM fructose did not significantly alter cell mass and volume in HBP. However, a statistically significant decrease of lipid dry mass was observed in 5mM fructose, suggesting channeling of carbons from fructose to other macromolecules other than lipids. Interestingly, the volume of lipids derived from 5mM fructose were not significantly distinct from those derived from 25mM glucose. This could be explained by the synthesis of different lipid species, supported by the observation that 5mM fructose resulted in lipid droplets of different size distributions, compared to 25mM glucose. These observations in HBP were consistent with MBP, where replacing glucose with fructose as the anabolic carbon source for adipogenesis resulted in lower levels of neutral lipid accumulation, as measured using fluorescently labeled lipid stains. Taken together, both glucose and fructose support differentiation to varying extents and the resultant fate of hexose-derived carbons are distinct. The use of stable isotope labeling with mass spectrometry may be able to uncover these differences, especially at the level of the specific lipid species favored by fructose vs. glucose.

Hexose combinations also influenced the nature of the lipids produced during HBP differentiation. Consistent with cell size changes, C4 and C5 yielded larger lipid mass and volumes, relative to C1. Again, the type of hexose present influences lipid synthesis more than the total amount of hexose carbons. When fructose is present with glucose, HBPs synthesize more lipids, and these lipids are more likely to be larger in diameter. These lipids also contribute more significantly total cell mass, accounting for > 30% of total cell mass in C5. To the best of our knowledge, these results represent the first study characterizing the physical properties of lipid droplets formed in the presence of different hexose sources. As lipid droplets consist of a central core of neutral lipids surrounded by a phospholipid monolayer containing integral and peripheral proteins^41^, it is possible that hexose combinations alter both lipid and protein composition in these organelles. Indeed, fructose has been described to promote de novo lipogenesis in hepatocytes^42^ while upregulating lipolytic genes in preadipocytes^43^. Thus, there is literature supporting the pleiotropic role of fructose in lipid remodeling, that might be responsible for altering lipid droplet size in BAT. In the same vein, fructose has also been shown to alter cellular proteostasis through ribosomal protein S6 kinase B1 (RPS6KB1) driving of hepatic protein synthesis^44^. While the same mechanisms have yet to be demonstrated in BAT, it is likely that hexose combinations coordinate both lipid and protein synthesis/degradation pathways, contributing to the lipid droplet heterogeneity seen in adipogenic differentiation.

There is emerging evidence that the fundamental physical properties of lipid droplets dictate cellular metabolism. In the liver, lipophagic internalization favors smaller lipid droplets, while larger lipid droplets are preferentially targeted for lipolysis^45^. Thus, lipid catabolism is in part driven by the physical size of these organelles. While our work has yet to uncover the consequences of fructose-specific alterations in lipid droplets, we surmise that cutting-edge, label-free imaging via optical diffraction tomography (ODT) is an ideal tool to study these metabolic changes in situ. In the future, combining non-destructive, optical imaging methods with spatially resolved matrix-assisted laser-desorption/ionization mass spectrometry (MALDI-MSI)^46^ is predicted to offer new biological insight in the relationship between cellular metabolism and physical dimensions of the cell.

## CONCLUSION

In this study, we employed biochemical and biophysical methods to assess the impact of hexoses on brown preadipocyte differentiation. We found that the availability of sugars influenced cellular hyperplasia and hypertrophy. In addition, optical diffraction tomography (ODT) was used to analyze human preadipocytes (HBP) during adipogenic differentiation under varying conditions of fructose, glucose, and a combination of both sugars. This approach is novel in its application of ODT to quantify key cellular metrics such as cell dry mass, cell volume, lipid dry mass, lipid volume, and lipid diameter throughout the differentiation process. Our findings reveal significant insights into how different sugars impact BAT morphology and lipid formation. We observed distinct changes in cell morphology and lipid formation in response to these conditions, highlighting the sensitivity of HBP to metabolic microenvironments.

## ACKNOWLEDGEMENTS

The authors would like to thank the Tseng Lab at Harvard Medical School (Boston, MA, USA) for the human brown preadipocyte cell lines.

Figures 2A and 3A were made with Biorender.com.

