## Supplemental Figure for "Sweet science: Exploring the impact of fructose and glucose on brown adipocyte differentiation using optical diffraction tomography"

**Supplementary Information**

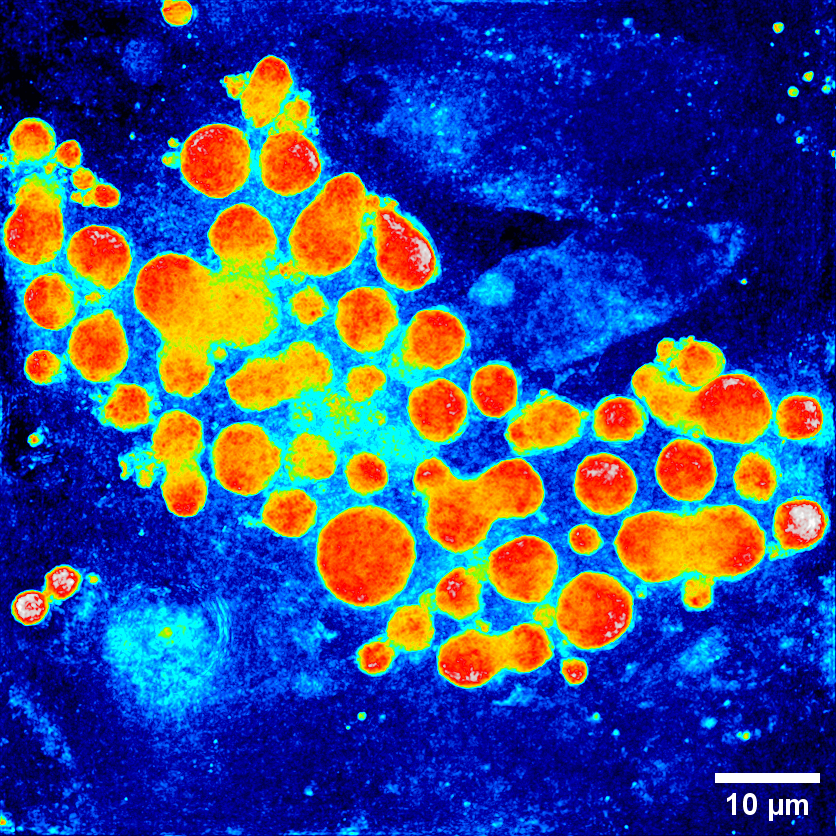

Figure S1: Example ODT image of a day 21, condition 5 HBP cell where fused lipid droplets are present.
